# CANCERSIGN: a user-friendly and robust tool for identification and classification of mutational signatures and patterns in cancer genomes

**DOI:** 10.1101/424960

**Authors:** Masroor Bayati, Hamid R. Rabiee, Mehrdad Mehrbod, Fatemeh Vafaee, Diako Ebrahimi, Alistair R.R. Forrest, Hamid Alinejad-Rokny

## Abstract

Analyses of large somatic mutation datasets, using advanced computational algorithms, have revealed at least 30 independent mutational signatures in tumor samples. These studies have been instrumental in identification and quantification of responsible endogenous and exogenous molecular processes in cancer. The quantitative approach used to deconvolute mutational signatures is becoming an integral part of cancer research. Therefore, development of a stand-alone tool with a user-friendly graphical interface for analysis of cancer mutational signatures is necessary. In this manuscript, we introduce CANCERSIGN as an open access bioinformatics tool that uses raw mutation data (BED files) as input, and identifies 3-mer and 5-mer mutational signatures. CANCERSIGN enables users to identify signatures within whole genome, whole exome or pooled samples. It can also identify signatures in specific regions of the genome (defined by user). Additionally, this tool enables users to perform clustering on tumor samples based on the raw mutation counts as well as using the proportion of mutational signatures in each sample. Using this tool, we analysed all the whole genome somatic mutation datasets profiled by the International Cancer Genome Consortium (ICGC) and identified a number of novel signatures. By examining signatures found in exonic and non-exonic regions of the genome using WGS and comparing this to signatures found in WES data we observe that WGS can identify additional non-exonic signatures that are enriched in the non-coding regions of the genome while the deeper sequencing of WES may help identify weak signatures that are otherwise missed in shallower WGS data.

## INTRODUCTION

Aberrant somatic changes in DNA resulting from endogenous sources (e.g. APOBEC-induced mutagenesis and DNA repair defects) and exogenous factors (e.g. tobacco smoking and UV radiation) are the hallmark of cancer. These alternations in DNA may have different forms, ranging from gross chromosomal rearrangements to single base substitutions [1]. The whole genome sequencing of tumor cells has shown that the number of mutations varies from less than one hundred per genome to hundreds of thousands depending on the cancer type and patient. Moreover, the type of mutation and sequence context of many cancer mutations are not random. For instance, C-to-T mutation within the CG (a.k.a. CpG) dinucleotide is a prevalent mutation in cancer and as its abundance is proportional to the age of patient it is referred to as an “aging” signature [2]. Many cancers also have a large number of C-to-T and C-to-G mutations within TCA and TCT trinucleotides [3]. These mutations are attributed to the aberrant changes in the level and activity of APOBEC enzymes. The mutational landscape of each cancer genome is thus a cumulative result of multiple mutational signatures, each caused by a unique process such as methylation, APOBEC mediated changes, etc. [1].

Typically, signatures of mutational processes are determined by considering the trinucleotide context of single base substitutions. If all mutations are presented based on changes in the same DNA strand, there are 96 possible different types of mutations within trinucleotide motifs [4]. In 2013, Alexandrov *et al*. proposed a mathematical framework for analysing mutational signatures [4] based on these 96 types of mutations. Using a matrix factorization algorithm, the authors uncovered 30 independent mutational signatures. Details of these signatures including their prevalence in each cancer type and potential etiology are available at the COSMIC database (http://cancer.sanger.ac.uk/cosmic/signatures [5]).

The discovery of mutational signatures was a breakthrough in the field of cancer research. Therefore, the mathematical framework developed by Alexandrov *et al*. [4] is now routinely used to identify novel mutational signatures and to study the processes involved in different cancers and in different patients. To help the progress of this field, we have developed a computational tool, CANCERSIGN, which enables the users to easily apply a matrix factorization analysis to cancer mutation datasets and receive a complete set of mutational signatures. Compared to the previously developed packages in R [6–8], CANCERSIGN is unique in that it is a stand-alone package (i.e. it does not require additional software programming). Therefore, to use this tool, no programming skills are required. Additionally, it enables the users to perform *de novo* mutational signature analyses.

Application of CANSERSIGN is not limited to extracting mutational signatures based on nucleotides immediately flanking the mutated site (i.e. tri-nucleotide motifs). It allows the users to extend the analysis to two bases on each side of the mutated base (i.e. penta-nucleotides motifs). According to a recent study [9], taking larger sequence contexts into consideration provides a greater power to explain variability in genomic substitution probabilities. In addition, CANCERSIGN allows the user to select trinucleotides of interest, and determine their penta-nucleotide mutational signatures. Furthermore, it has a built-in clustering option to study the groupings of cancer samples based on the raw mutation counts and/or composition of mutational signatures. In this manuscript, we introduce CANCERSIGN and show the new mutational signatures obtained from a *de novo* analysis of whole genome ICGC dataset. This analysis was performed for each cancer type separately and resulted in 77 mutational signatures. Each of the obtained signatures were shown to be highly similar to at least one of the 30 signatures discovered previously [2], except five signatures that can be considered as novel.

## DATA and METHODS

To develop CANCERSIGN and demonstrate its application, we used all the whole genome mutation datasets available at the International Cancer Genome Consortium (ICGC) data portal. A summary of the datasets used is shown in **Supplementary table 1**.

A schematic of CANCERSIGN features is given in **Supplementary figure 1**. This tool deciphers mutational signatures in somatic mutation datasets using a previously reported non-negative matrix factorization (NMF) model [4]. It only requires a dataset of mutations in a simple format described in the tool manual (https://github.com/masroor-bayati/cancer_analysis_tool and https://github.com/ictic-bioinformatics/CANCERSIGN).

CANCERSIGN then performs the requested analyses and outputs mutational signatures (3-mer and/or 5-mer formats) and clustering figures. The numerical outputs of the analyses are saved in a folder named “output” (refer to the CANCERSIGN manual). This data can be used to gain further insight into mutational processes in cancer. For example, the exposure matrix obtained from the NMF analysis, can be used to determine the prevalence of each mutational signature in each sample.

### Estimation of mutational signatures

Each mutational signature is defined by a total of 96 types of mutations within tri-nucleotide motifs [4]. Assuming that 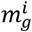 represents the number of mutations of type *i* recorded in the mutational catalogue of sample *g*, then the mutational catalogue of multiple samples is represented by a matrix *M* (*K* × *G*):

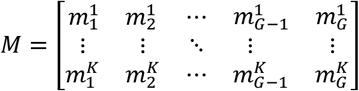

Where, *K* represents the number of mutation types (K=96 for tri-nucleotide signature analysis) and *G* represents the number of cancer samples. Therefore, the ίth column of *M* has the number of each of 96 mutation types in the ίth sample. Here, it is assumed that the mutational catalogue of each sample (i.e. each column of M) is the result of linear superposition of several mutational signatures [4], each of which corresponds to a particular mutational process. Assuming that the number of mutational signatures is N, we need to factorize *M* into two matrices *P* and *E* with sizes *K* × *N* and *N* × *G*, respectively:

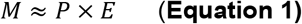

The expanded representation of the above equation is given by:

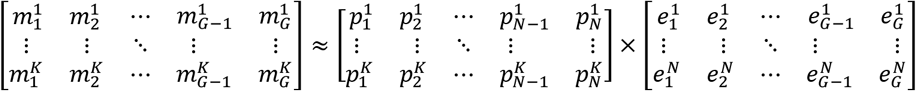

Here, each column of *P* is interpreted as one mutational signature and each row of *E* represents the exposures of each mutational signature in the corresponding sample genome (i.e. the prevalence of each mutational signature in that sample). The process of estimating independent mutational signatures from a mutational catalogue matrix is done using a nonnegative matrix factorization (NMF) method. Note that the elements of the input catalogue matrix are nonnegative because they represent the number of mutations.

The algorithm used in this framework consists of several iterative steps. The overall procedure is as follows. Each iteration starts with sampling from matrix *M* and creating a new bootstrapped matrix 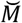. To do this, each column of *M* (mutation counts of one input cancer genome) is considered a discrete probability distribution and is resampled to create the corresponding column of 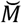. Next, the bootstrapped matrix 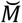 is factorized into matrices *P* and *E* by applying the NMF algorithm. These steps will be repeated for a certain number of iterations (typically 600 iterations are enough to obtain a stable result). The resulting matrices obtained after each iteration are stored and grouped into two sets: *P* matrices and *E* matrices named *S_P_* and *S_E_*, respectively. The set of columns of matrices in *S_P_* are then clustered into *N*groups using a variation of the *k - means* clustering algorithm [10]. A similarity measure between two columns is calculated using cosine similarity and the centroids of the clusters are obtained by averaging over the members of each cluster. The cosine similarity between the vectors *A* and *B* is calculated as 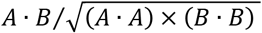, where *A · B* represents the dot product of vectors *A* and *B*. Finally, the *N* centroids are combined to form the columns of a single matrix 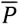, which is the matrix of mutational signatures. Since each column of matrix *P* corresponds to one specific row of matrix E, the procedure of clustering columns of matrices in *S_P_* naturally leads to clustering of rows of matrices in *S_E_*. Therefore, the final exposure matrix 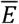 (prevalence of mutational signatures) is constructed in a manner similar to 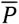 after the clustering step. Finally, the results are evaluated by calculating the reproducibility of mutational signatures and the Frobenius reconstruction error of the NMF solution. The reproducibility is measured by calculating the average silhouette width of the result of clustering step, while the reconstruction error of the solution is quantified by the Frobenius norm of difference between matrix M, and its estimation obtained by the factorization algorithm, i.e., 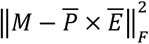 [4]. The overall procedure described above is carried out for different values of N; the number of deciphered signatures. The range of values for *N* can be specified by the user and the optimum value of *N* can be selected based on the aforementioned evaluation measures [4]. An optimum *N* leads to high reproducibility and low reconstruction error. CANCERSIGN produces an evaluation plot similar to the one proposed by Alexandrov *et al*. [4].

CANCERSIGN is designed to carry out each iteration of the aforementioned algorithm, which consists of one bootstrap and a complete NMF decomposition, independent and in parallel with other iterations. This enables the user to distribute the whole task across CPU cores and speed up the procedure of extracting mutational signatures. This tool also allows the user to specify the convergence criteria, the number of NMF iterations, the number of bootstrapping iterations and the number of CPU cores for parallelization.

### Penta-nucleotide mutational signatures

In addition to extracting mutational signatures based on the immediately flanking nucleotides around the mutated site (tri-nucleotides), CANCERSIGN enables the user to further investigate the underlying mutational process, by expanding the mutational context to two bases upstream and downstream of the mutated base (penta-nucleotides). For this purpose, the user can choose an arbitrary set of mutations in tri-nucleotide motifs (up to 10 combinations). For each of these selected motifs, our tool quantifies the number of mutations in the corresponding 16 penta-nucleotide motifs (e.g. when the motif C[G>T]A is selected, 16 penta-nucleotide motifs of the form NC[G>T]AN are considered where N is each of four nucleotides). The mutation counts of penta-nucleotides in all samples are then used as the input to the NMF analysis, which produces a set of penta-nucleotide (or 5-mer) mutational signatures.

### Clustering of samples

One important application of CANCERSIGN is clustering of tumors based on mutations within selected motifs (Matrix *M* in eq. 1) or based on the prevalence of mutational signatures (Exposure Matrix *E* in eq. 1). This feature allows the user to investigate between sample heterogeneity and to identify potential outlier samples. Additionally, this information can be used to investigate correlation with clinical data including treatment history and survival outcomes.

The clustering of samples is done using the k-means algorithm with Euclidian distance measure. For this purpose, the NbClust function from NbClust package [11] in R is utilized which obtains the optimal number of clusters automatically based on 30 indices. To visualize the result of clustering, CANCERSIGN produces two types of plots. The first plot visualizes the clustering centroids, which are obtained by averaging over features of the members of each cluster. The second plot visualizes a projection of the samples in a 2D plane to show how samples within a cluster are similar to each other and different from samples in other clusters. To do this, the tool performs a principal component analysis (PCA) and generates a PC1-PC2 plot. The user can find the sample IDs associated with each cluster by using tabular files produced by the tool (see CANCERSIGN manual). A snapshot of CANCERSIGN is provided in Figure 1.

**Figure 1.**
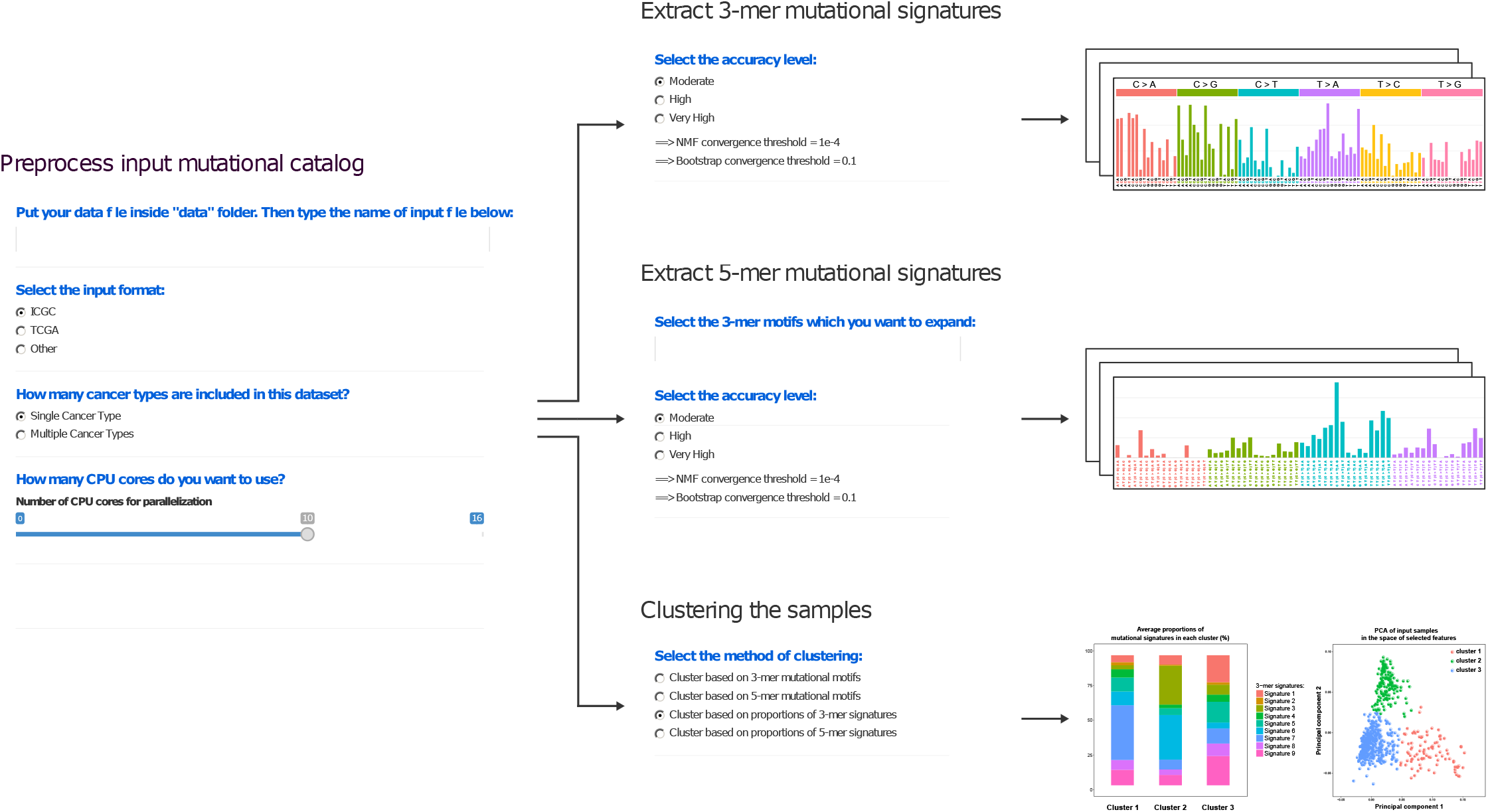
Tool functionality map. This diagram abstracts the functional features of our tool. The user can either choose to extract mutational signatures in the conventional format which are the spectrums over 96 mutation types (3-mer motifs) or extract 5-mer mutational signatures. The box indicating “NMF algorithm” consists of several steps. Its main steps are bootstrapping from mutational catalogue matrix, NMF iteration and clustering the results obtained from the bootstrapped data. In the clustering part, the user can choose different sets of features to cluster the samples. These feature sets are based on either mutational motifs or mutational signatures. In the former case, counts or proportions of 3-mer or 5-mer mutational motifs can be selected, and in the latter case, contributions of 3-mer or 5-mer mutational signatures can be chosen for clustering features. Finally, in the simulation part, there are two options to simulate the mutational catalogues and in either case, it produces simulated signatures to examine the statistical significance of deciphered signatures from the input data.

We have applied our tool to the mutational data of various cancers (18 types-**Supplementary table 1**) and presented the results. The ICGC somatic mutations dataset (last updated June 2017) is used for this analysis. General information and statistics about the data can be found in **Supplementary table1**. Details about our analyses, including the tool parameters, are provided in supplementary information.

## RESULTS AND DISCUSSION

The results of separate analyses of whole genome mutation data from each of 18 tumor types reported in the ICGC database are presented in Figures 2A-2R. The evaluation diagrams for each tumor type are presented in **Supplementary figures 2-19**. These diagrams were used to determine the total number of mutational signatures presented in Figure 2. For each tumor type, the numerical values of the deciphered 3-mer signatures and the prevalence of each signature in each tumor sample are obtained and provided in **Supplementary tables 2-19**. Totally, The CANCERSIGN program identified 77 signatures in 18 tumor-types (Figures 2A-2R). The majority of these signatures were highly similar (Cosine Similarity > 50) to one or more of the previously reported 30 signatures [2]. But five of these signatures had a Cosine similarity of <50, suggesting that they are possibly novel signatures that have not been identified previously (Figure 3). For example, Figure 2A indicates nine mutational signatures deciphered from the mutational data of breast cancer samples by using CANCERSIGN. For each of these nine signatures there is at least one previously reported signature which can be considered as a close match (Figure 3). Figure 2B shows the mutational signatures that CANCERSIGN has identified in blood cancer samples. For this cancer type, one signature (Blood Signature No. 1) does not resemble any of the previously reported signatures, i.e. it does not have >50% similarity to any of the previously reported signatures (Figure 2B). Details of the correlations between all 77 signatures discovered by CANCERSIGN, and the signatures reported by Alexandrov *et al*. [2] are given in **Supplementary table 20**.

**Figure 2.**
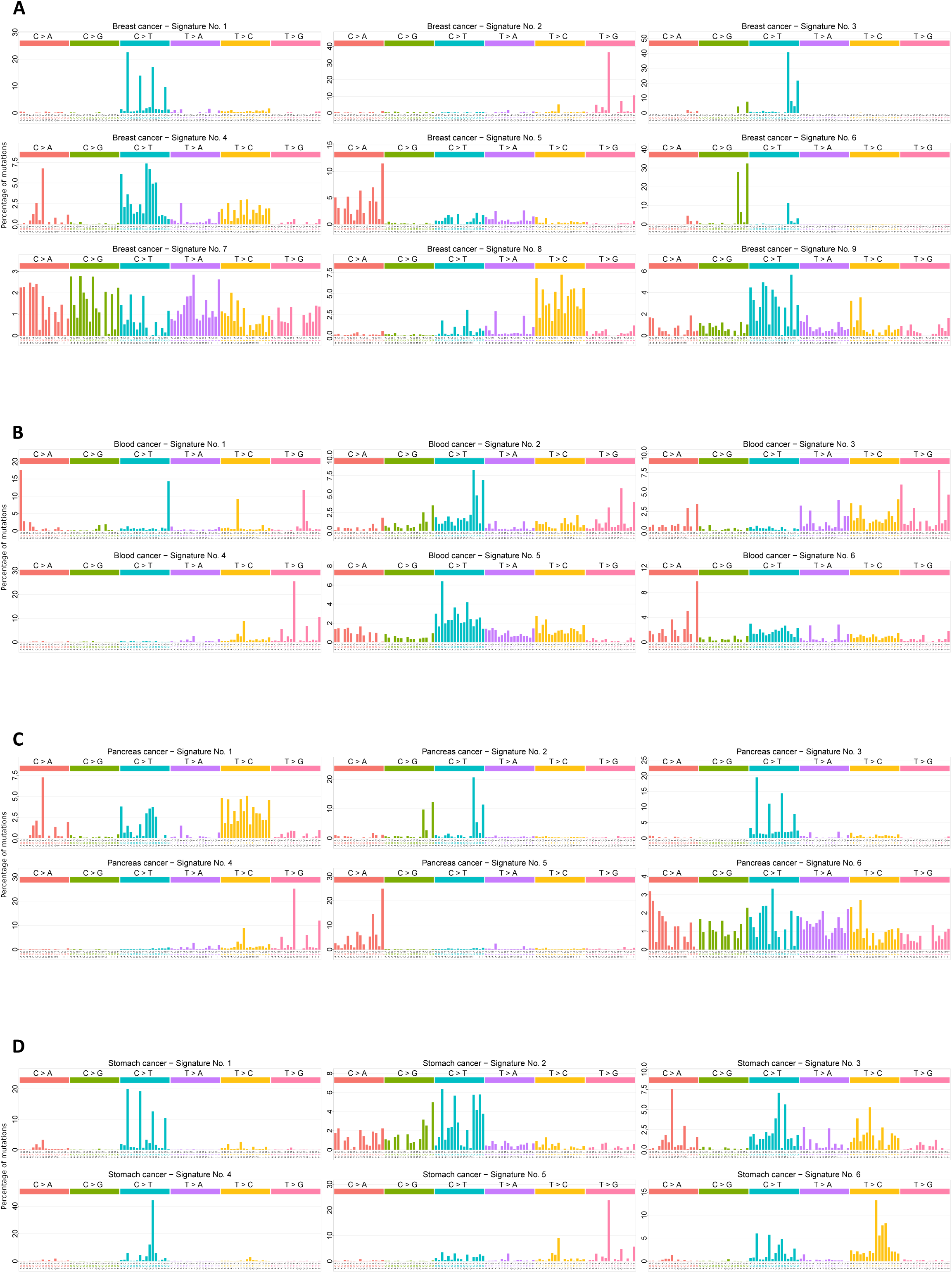
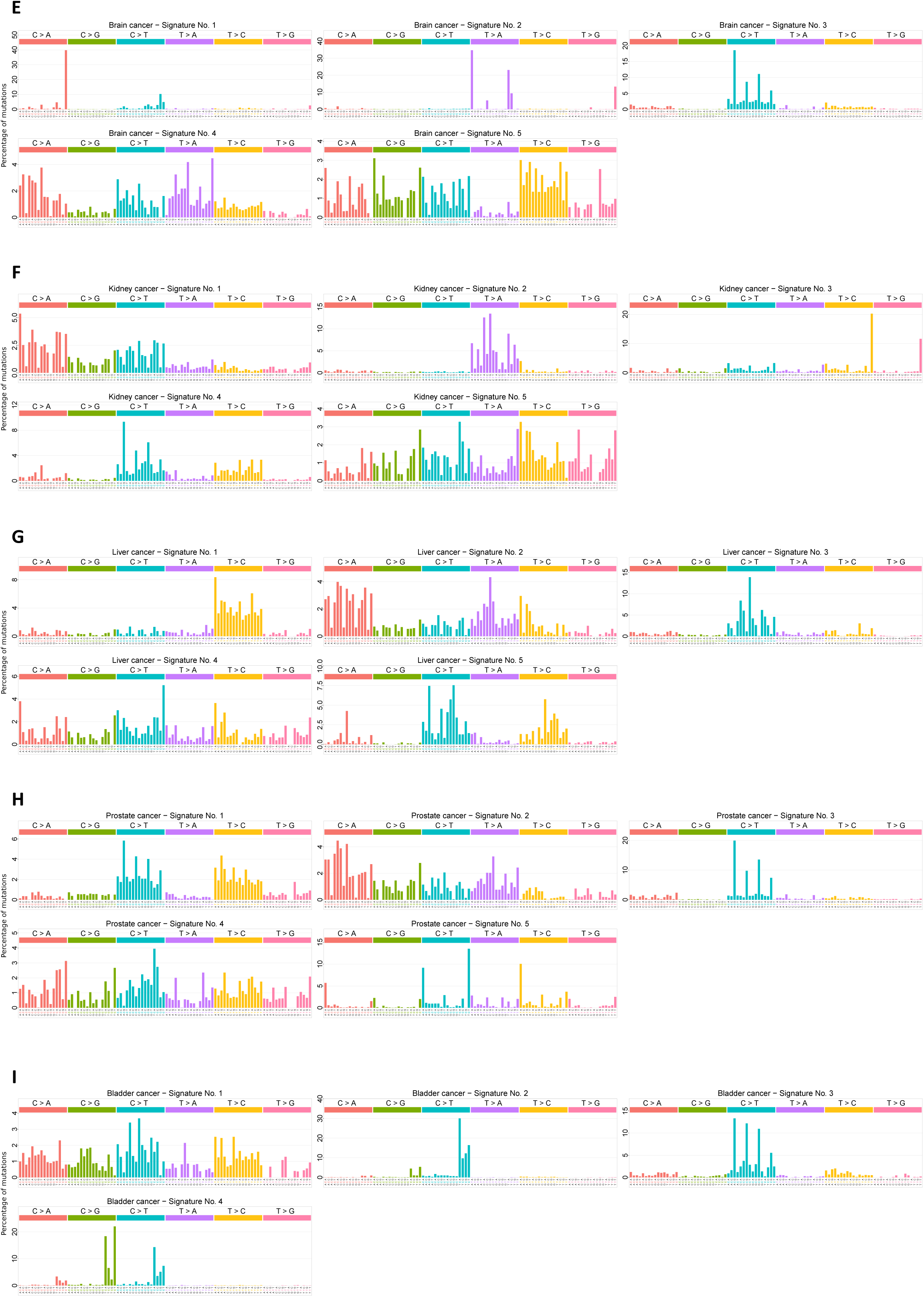
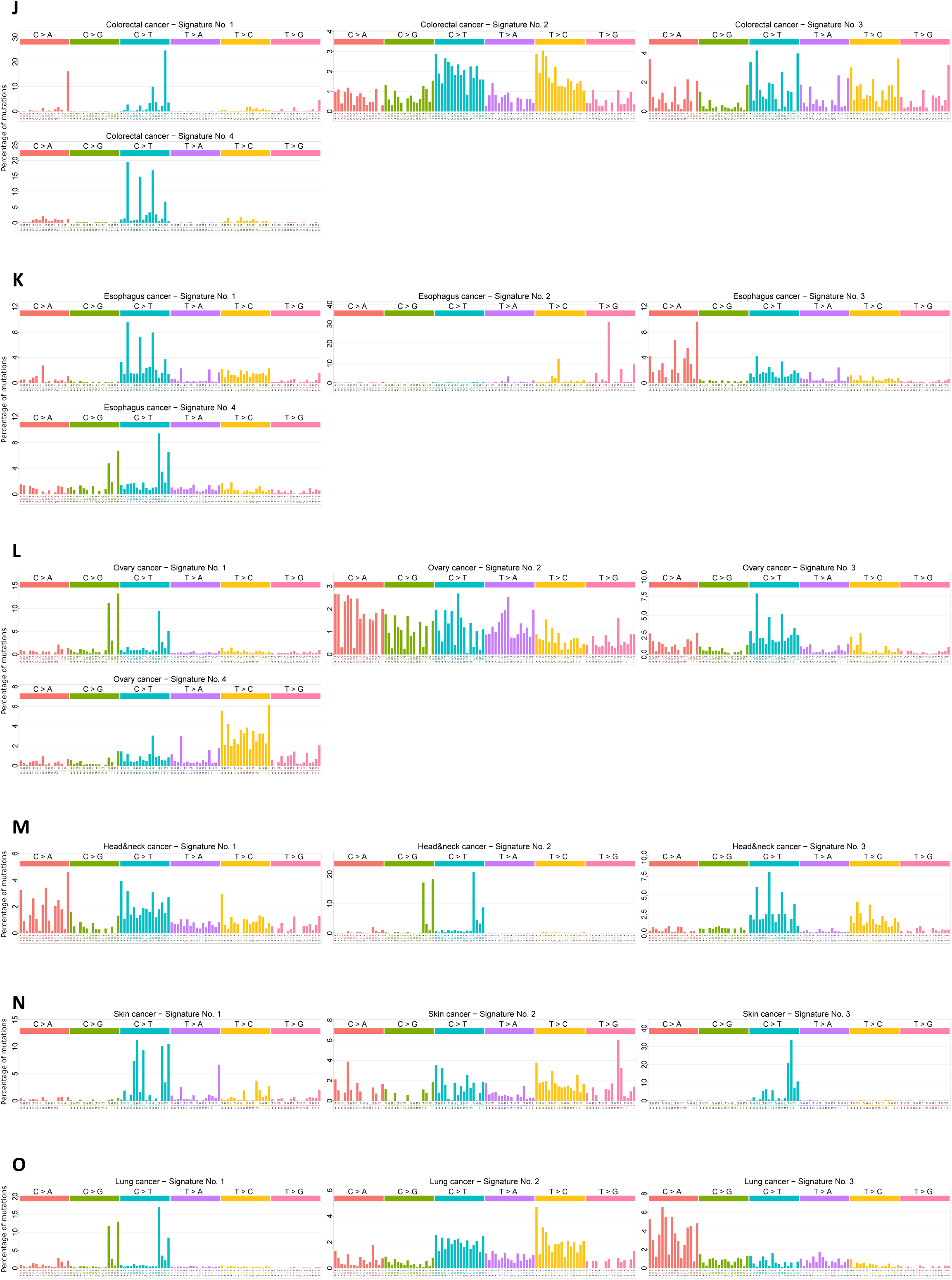
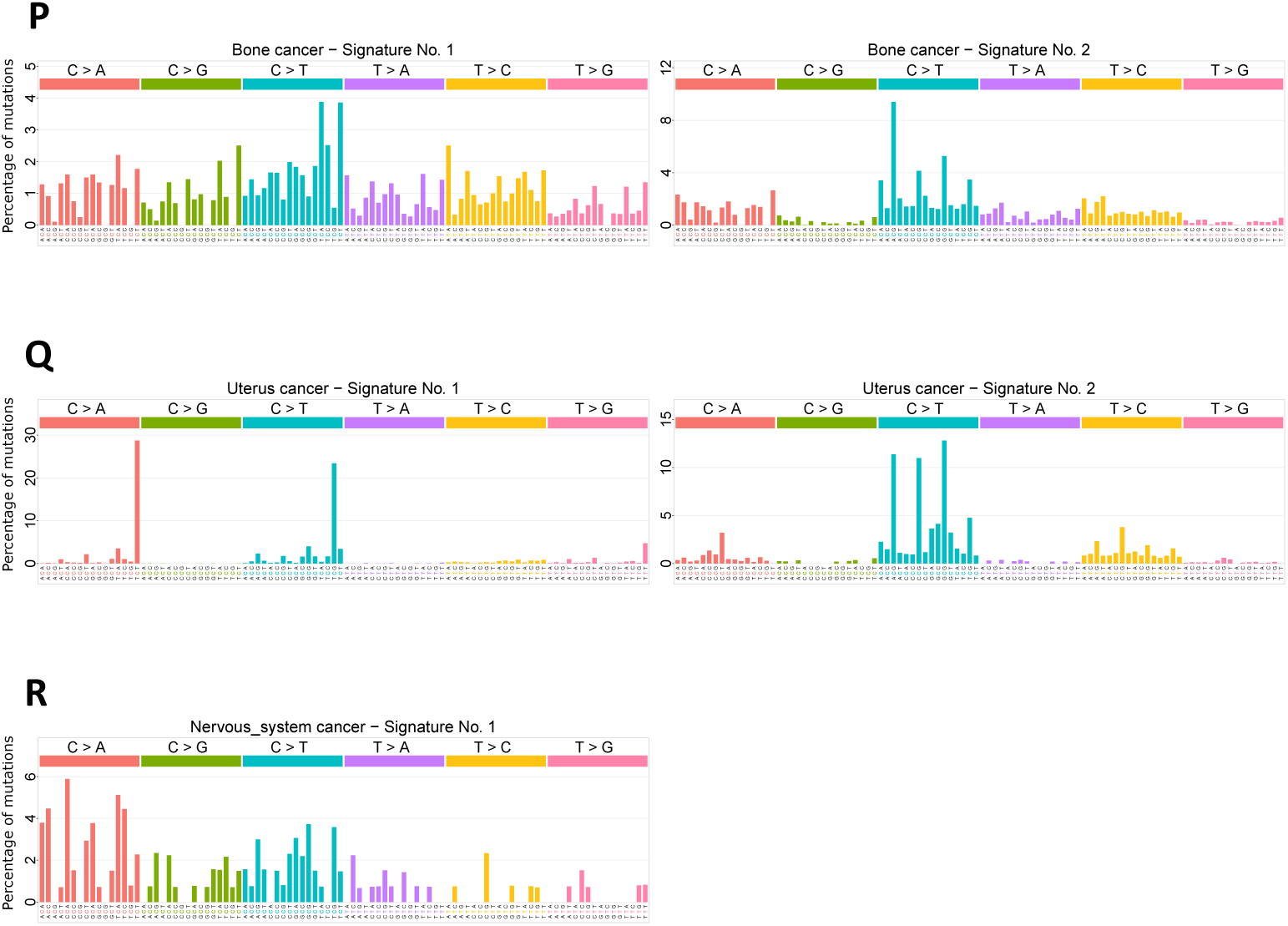
Deciphering whole genome 3-mer mutational signatures for various cancer types. The 3-mer mutational signatures extracted from whole-genome samples of 18 cancer types. Each cancer type is analysed separately. (A) Breast cancer 3-mer signatures. (B) Brain cancer 3-mer signatures. (C) Esophagus cancer 3-mer signatures…(R) Nervous system cancer 3-mer signatures.

**Figure 3.**
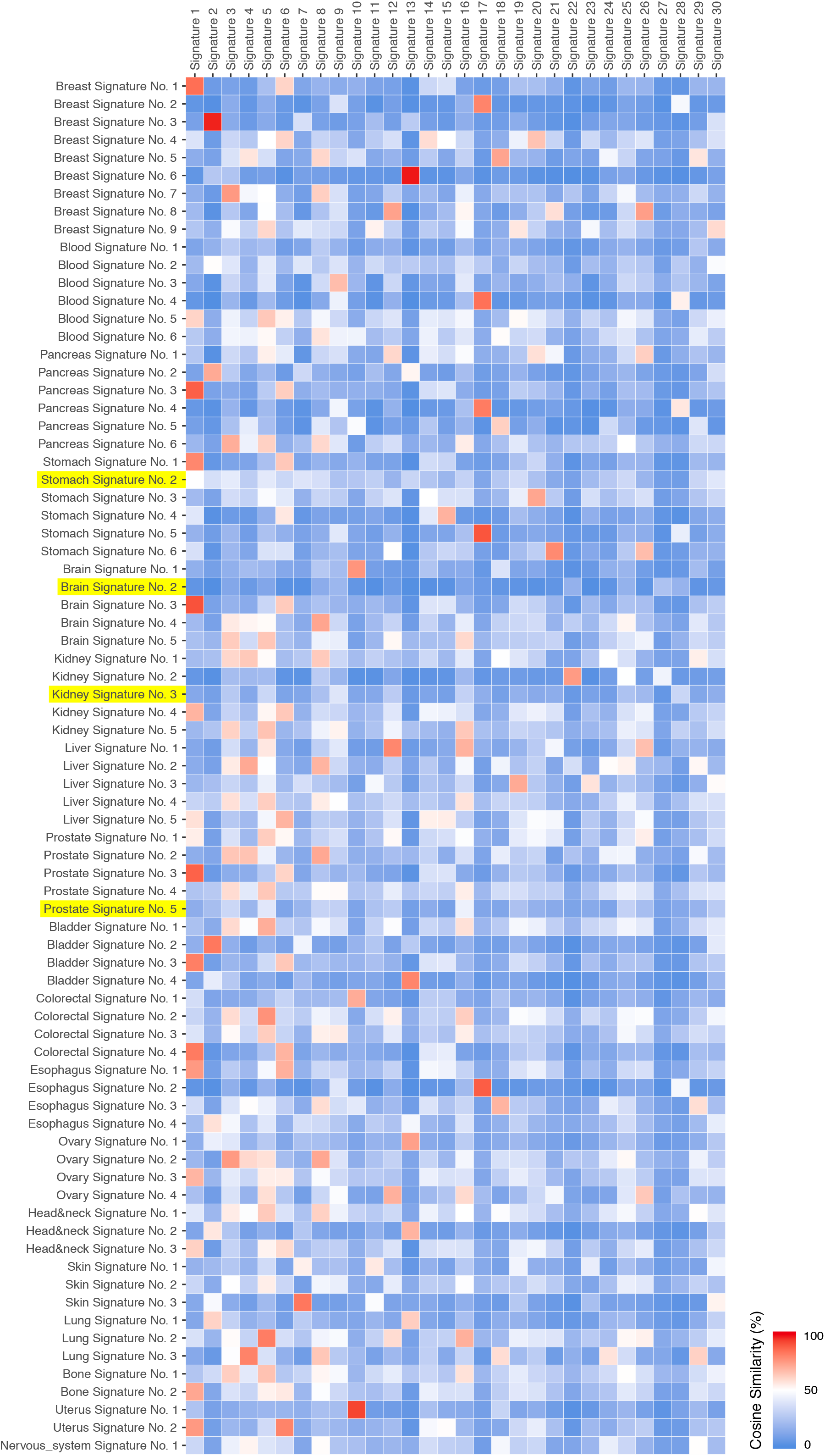
Correlation of 77 signatures identified by CANCERSIGN with 30 previously reported signatures by Nik-Zainal et al. [15]. Here, each cell indicates percentage of similarity between signatures identified by our tool and previously reported signatures. Those signatures with less than 50% similarity with previously reported signatures have been highlighted.

The 30 mutational signatures reported previously, is based on the analysis of all tumor types together. These signatures seem to be a consensus and represent trends across all tumor types. However, this global mutational signature analysis of multiple cancer types might produce results that are biased toward signatures of the tumors with more samples in the dataset. Furthermore, pooling somatic mutations obtained from whole exome sequencing with that of whole genome data can be another source of bias that is not considered in the previous studies. These issues have been discussed in a recent review by Nik-Zainal and Morganella [12]. Nevertheless, our tool can be used to extract mutational signatures for all tumors together, or for each tumor separately. Typically, researchers use mixed datasets (whole genome + whole exome) to extract mutational signatures. CANCERSIGN is able to provide an option for users to extract mutational signatures from the whole genome, whole exome or pooled data. For example, Figures 4 and 5 show the analysis of whole exome alone or whole exome + whole genome for breast and skin cancers, respectively (The evaluation diagrams for these analyses are presented in **Supplementary figures 20-23**). Figures 2A and 2N also show the analyses on whole genome data for breast and skin cancers, respectively. Here, we identified ten signatures in breast cancer pooled data (WGS+WXS) analysis. In the breast cancer whole genome analysis, we identified nine signatures; the signature No. 7 in Figure 4A disappeared from the whole genome data analysis and seems to be a WXS-specific signature. We then used breast cancer whole exome data as an input for CANCERSIGN and identified five signatures, in which one of them (Signature No. 1 in Figure 4B) has not been appeared in either pooled or whole genome data analyses. As a result, there are two signatures (Signature No. 1 and Signature No. 4 in Figure 4B) in our analyses that seems to be associated with only whole exome data. To find out if this is a sequencing bias, we extracted exome parts of whole genome breast cancer samples; here, CANCERSIGN identified three signatures (Figure 6) that are identical with three of the five whole exome analysis signatures (Signatures No. 2, No. 3, and No. 5 in Figure 4B). Signatures No. 1 and No.3 (Figure 4B) did not appear in this analysis, suggesting that these two signatures are specific for whole exome samples. Our detailed investigation of the breast cancer whole exome samples revealed that these samples have been sequenced ~2.5x more deeply than whole genome samples. This may indicate that the two WXS-specific signatures are rare signatures and can be seen in the deeply sequenced samples. We then extracted non-coding regions of the WGS breast cancer samples and used them as an input for CANCERSIGN. This analysis revealed nine signatures as the same with the whole genome data (Figure 6); interestingly, five of the nine signatures did not appear in the analysis of the WXS samples or exonic regions of the WGS samples, suggesting that these five signatures are non-coding region specific.

**Figure 4.**
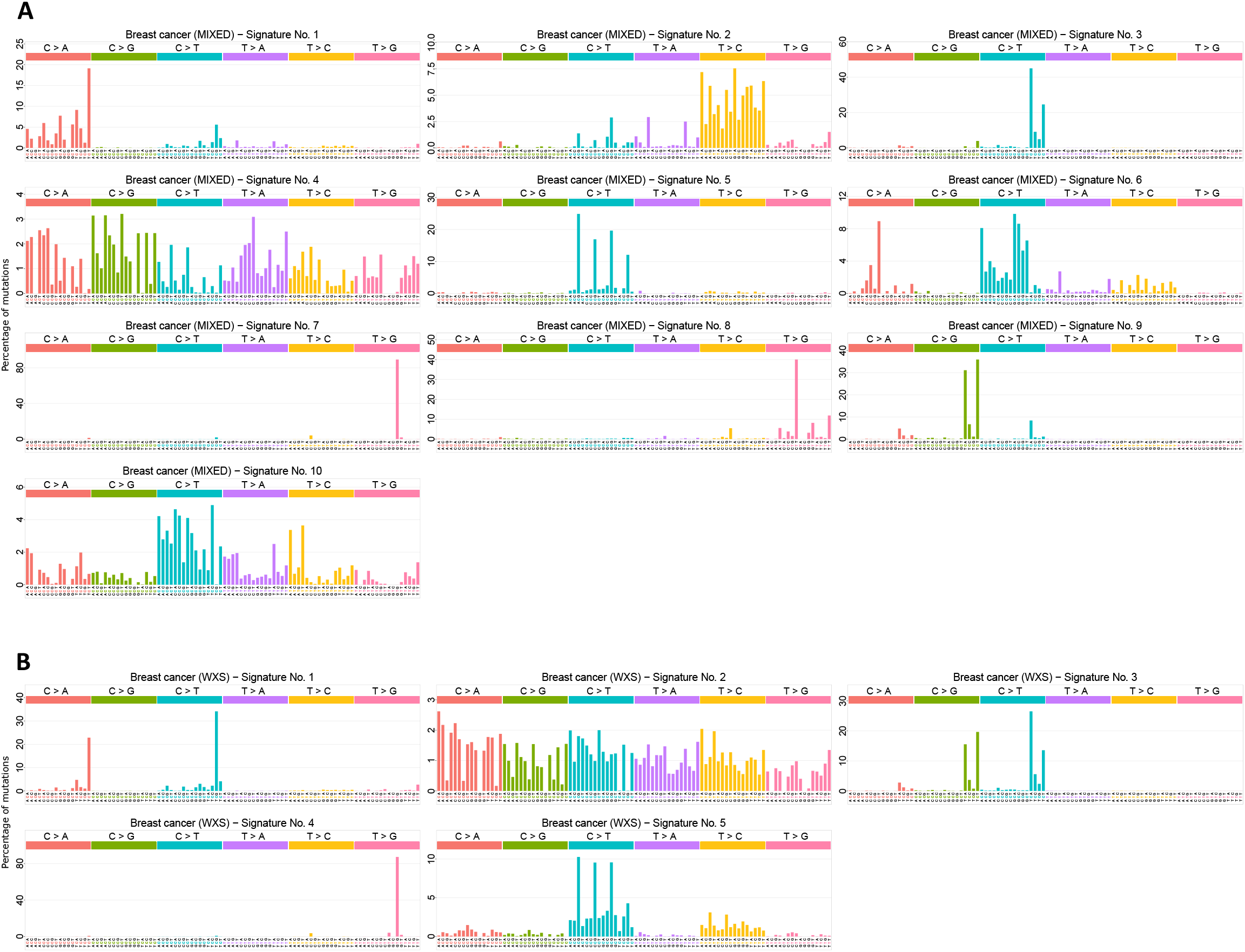
Mutational signatures deciphered from the whole exome and pooled (whole genome and whole exome) data of breast cancer. (**A**) For the pooled data. (**B**) For the whole exome data.

**Figure 5.**
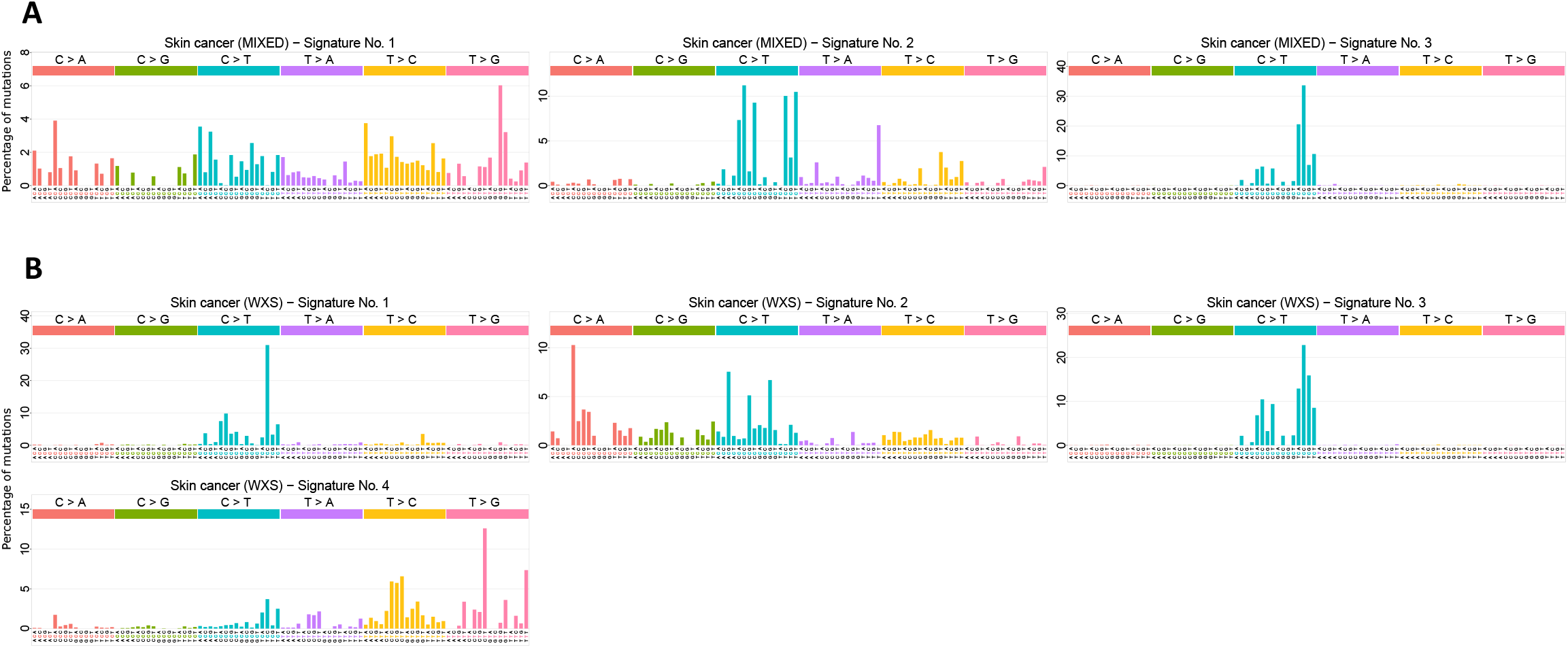
Mutational signatures deciphered from the whole exome and pooled (whole genome and whole exome) data of skin cancer. (**A**) For the pooled data. (**B**) For the whole exome data.

**Figure 6.**
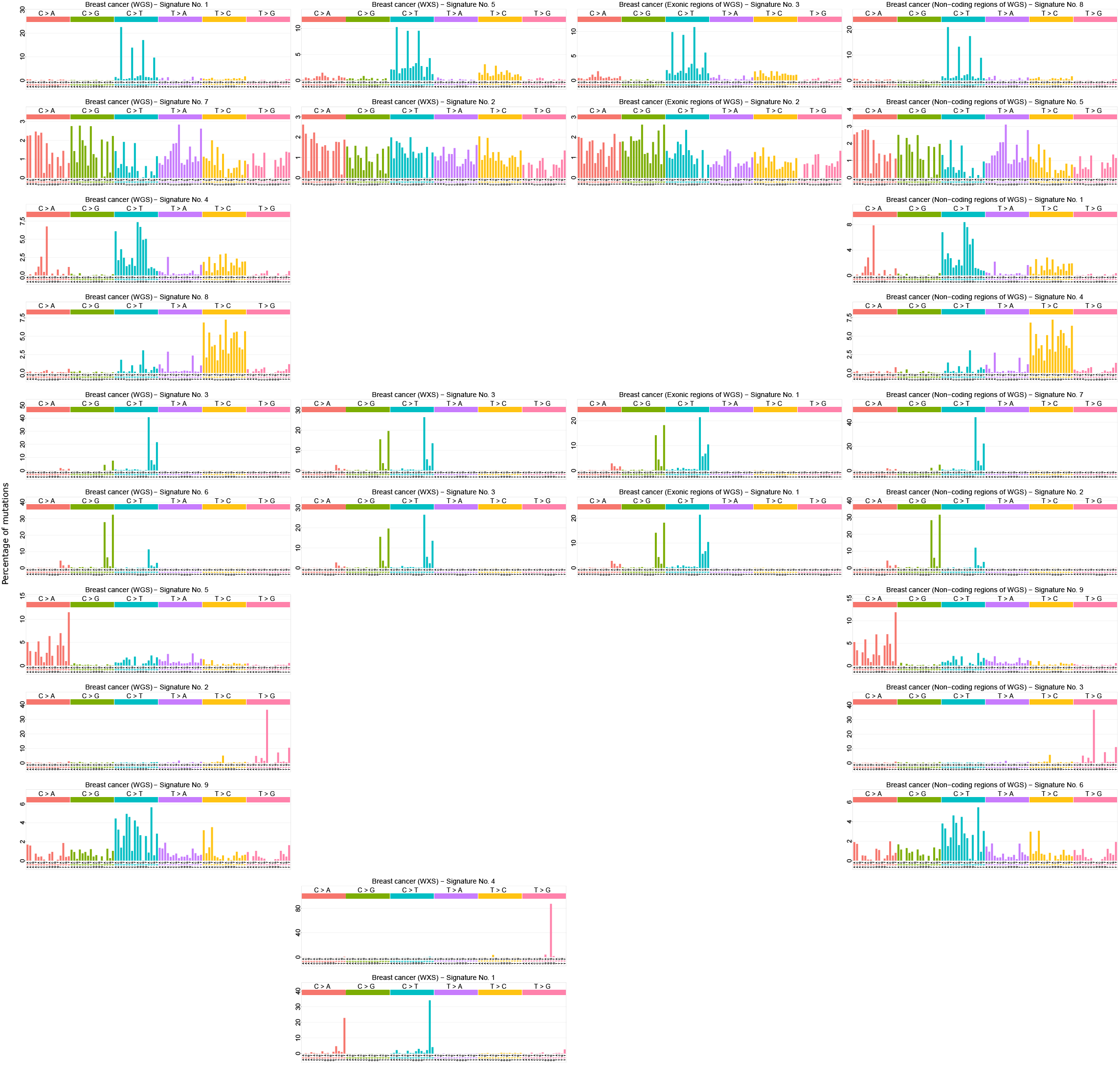
Mutational signatures deciphered from the whole genome, whole exome, exonic section of whole genome and non-coding regions of whole genome breast cancer samples. (**A**) Nine signatures discovered from whole genome analysis of breast cancer sample by CNCERSIGN. (**B**) In the analysis of whole exome sequencing breast cancer samples, CANCERSIGN detected five signatures. (**C**) CANCERSIGN also analysed exonic sections of WGS breast cancer samples and identified three signatures. (**D**) In the analysis of non-coding regions of the WGS breast cancer samples, CANCERSIGN detected nine signatures, in which five of them are seems to be non-coding specific signatures.

Identification of differences between tumor samples using clustering feature of CANCERSIGN can be informative, because this output can be used along with clinical data such as subtype, response to therapies, and overall survival to identify potential associations. To demonstrate the clustering feature of CANCERSIGN, we used breast cancer samples and performed both of the clustering options available in this tool. One option is clustering based on the proportion of mutational signatures. The results of this clustering is shown in Figure 7. Based on the estimated proportion of the nine signatures of breast cancer, CANCERSIGN classified the samples into three distinct groups (Figure 7). The second option is clustering based on the proportions of 3-mer motifs. The selected motifs were T(C->G)A, T(C->G)T, T(C->T)A and T(C->T)T. The result of this clustering is shown in Figure 8.

**Figure 7.**
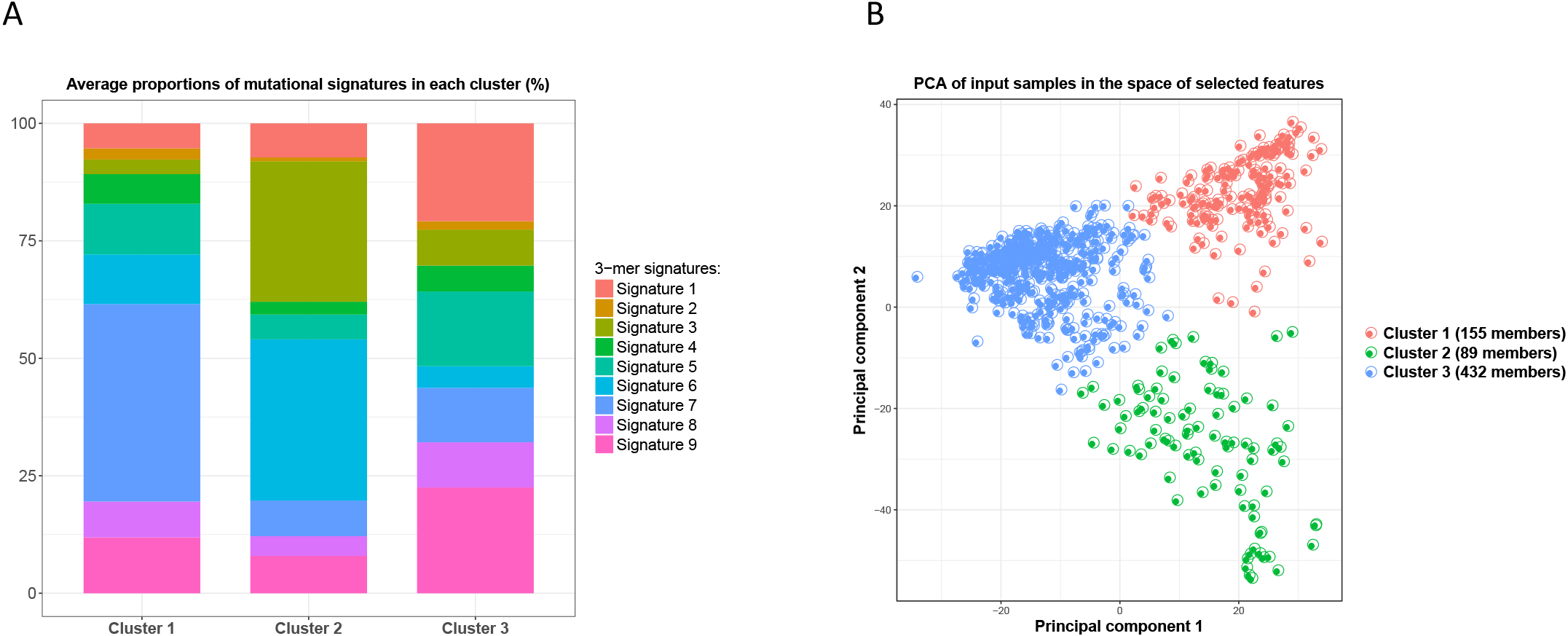
Clustering the samples of breast cancer based on contributions of 3-mer mutational signatures. This clustering is applied to whole-genome samples of breast cancer. The features of samples for this clustering are the contributions of 3-mer mutational signatures to each sample. (**A**) Average contributions of signatures within each cluster. (**B**) The result of principal component analysis of all samples in the space of clustering features. The axes of the plot are the first two principal components.

**Figure 8.**
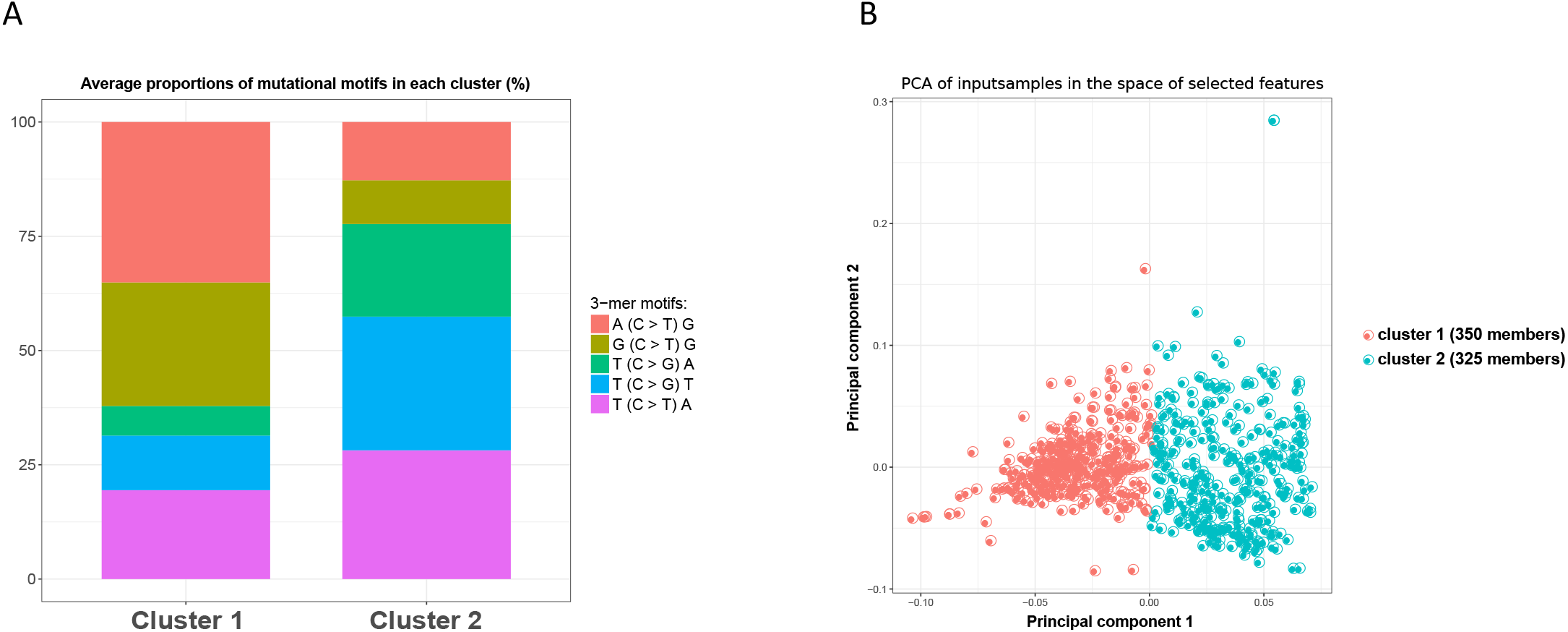
Clustering the samples of breast cancer based on proportions of 3-mer mutational motifs. This clustering was applied to whole-genome samples of breast cancer. The features of samples for this clustering are proportions of the 3-mer mutational motifs recorded in each sample. (**A**) Average counts of motifs within each cluster. (**B**) The result of principal component analysis of all samples in the space of clustering features. The axes of the plot are the first two principal components.

Another feature of CANCERSIGN is to decipher mutational signatures based on mutations within specific 5-mer motifs. To demonstrate this feature, we selected two 3-mer motifs (TCA and TCT) and two mutation types (C>T and C>G) as an input (i.e. T(C>G)A, T(C>G)T, T(C>T)A and T(C>T)T). These mutations are attributed to APOBEC enzymes and are the most abundant mutations in many cancer types [13] [14]. The CANSERSIGN program quantified the number of mutations within all possible 5-mer motifs that have been obtained from these 3-mer motifs. As an example, for C>G mutation within the 3-mer TCA motif, the program correctly counts the number of mutations for all 16 possible 5-mers (i.e. NT(C>G)AN where N:A, C, G, or T). The analysis by CANCERSIGN, over all the selected mutations in breast cancer samples, revealed four 5-mer mutational signatures, as shown in Figure 9. The evaluation diagram for this analysis is given in Supplementary figure 24. The numerical values of the resultant 5-mer signatures and their prevalence for each breast cancer sample are provided in **Supplementary table 21**.

**Figure 9.**
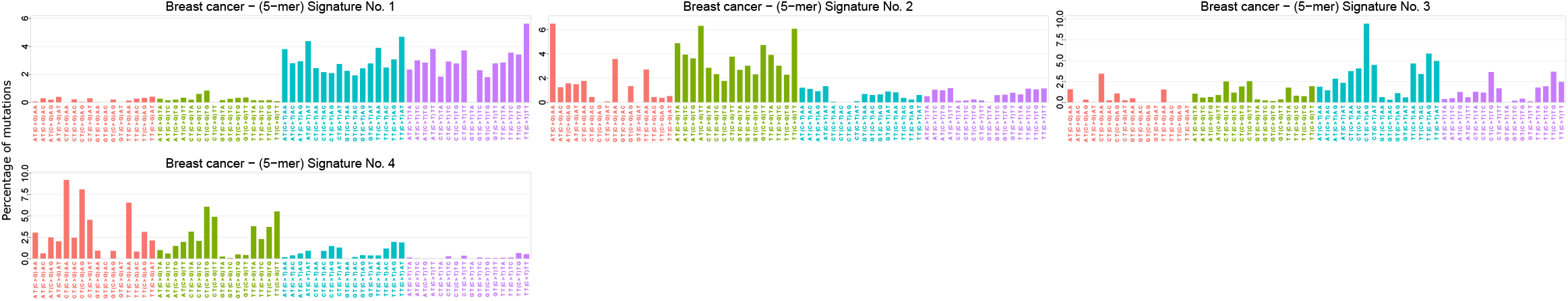
Deciphering whole genome 5-mer mutational signatures for breast cancer. The 5-mer mutational signatures extracted from whole-genome samples of breast cancer. Here, we have chosen four 3-mer motifs, namely T(C>G)A, T(C>G)T, T(C > T)A and T(C > T)T, to expand to 5-mer motifs and extracted the corresponding 5-mer signatures.

### Comparison with other tools

We have compared our tool with its well-known counterparts which have been developed for similar purposes: SomaticSignatures [6], SigneR [7] and deconstructSigs [8]. The deconstructSigs package is different from other tools as it is designed to determine the optimal linear combination of pre-defined mutational signatures that most accurately reconstructs the mutational profile of a single tumor sample [8], whereas others are designed to extract *de novo* mutational signatures from a cohort. Table 1 compares the tools based on a set of features. Note that this comparison only considers relevant features of these tools rather than all capabilities. Please refer to the corresponding papers for more information [6–8].

**Table 1.** A comparison between packages for analysis of mutational catalogues.

|  | Usage | Method of selecting optimum number of extracted signatures | Option of extracting 5-mer mutational signatures | Option of clustering the samples based on motifs or signatures | Graphical User Interface | Implementation |
| --- | --- | --- | --- | --- | --- | --- |
| **CANSERSIGN** | Deciphering *de novo* mutational signatures | <u>Specified by user by</u> inspecting the diagrams of reproducibility and reconstruction error | Yes | Yes | Yes | R |
| **SomaticSignature** | Deciphering *de novo* mutational signatures | <u>Specified by user by</u> inspecting the diagrams of residual sum of squares and explained variance | No | No | No | R |
| **SigneR** | Deciphering *de novo* mutational signatures | <u>Automatically</u> specified by empirical Bayesian treatment to the Poissonian NMF model | No | No | No | R |
| **deconstructSigs** | Obtain the optimal linear combination of pre-defined mutational signatures | — | No | No | No | R |

The tools were applied to a simulated dataset of mutations. We constructed the simulated dataset as follows: 6 signatures were selected from the list of mutational signatures presented at the COSMIC database [5] (signatures 1, 2, 3, 17, 21 and 29) as the underlying mutational processes, producing our simulated mutational catalogue. The mutational load (total mutations) for each sample was determined by taking a random number from a Rayleigh distribution with parameter *σ* = 8000 (this distribution was used since we observed that the histogram of mutational loads for samples of a cancer is often unimodal and positively skewed, and the average of mutational loads across all samples and all cancer types is near 10’000 which is approximately equal to the mean of a Rayleigh distribution with scale parameter of 8000). In the next step, the vector of mutations for each sample was generated by a linear combination of the 6 signatures where the 6 coefficients were determined randomly (for each set of coefficients, we selected 6 random numbers between 0 to 1, then normalized them such that they sum up to 1, and finally multiply them with the amount of mutational load of the sample). The aforementioned algorithm were used to generate 400 samples to form the simulated mutational catalogue.

We tried to apply CANCERSIGN, SomaticSignatures and SigneR to the simulated dataset in an equal condition. However, due to its Bayesian framework, SigneR is not scalable to large mutational catalogues containing several hundreds of samples. Consequently, it is not feasible to test SigneR on the simulation dataset, and instead performed the comparison between CANCERSIGN and SomaticSignatures.

The parameters of the analysis were set as follows. The range of values of ***N*** (number of signatures to decipher) was set from 2 to 12. The maximum number of bootstraps for each ***N*** was set to 100 for CANCERSIGN, and the number of replicates (nReplicates) for SomaticSignatures was set to 20. With these settings, the tools consumed approximately the same amount of time (~45 minutes) to decipher mutational signatures from our simulated dataset (using a typical computer with four 1.7 GHz CPU cores and 8GB memory). According to Figure 10, both tools have correctly found *N* = 6 as the optimal number of underlying mutational signatures (the knee point in the diagram of summary statistics of SomaticSignatures [6], and the point with a high reproducibility and the lowest reconstruction error in the evaluation diagram of CANCERSIGN). The obtained mutational signatures are shown in Figure 11. By a simple visual comparison, we can conclude that both tools have deciphered almost identical signatures which are also identical to the original selected signatures (signatures 1, 2, 3, 17, 21 and 29 from the COSMIC database). This test shows that both tools can produce valid results when analysing catalogues of somatic signatures.

**Figure 10.**
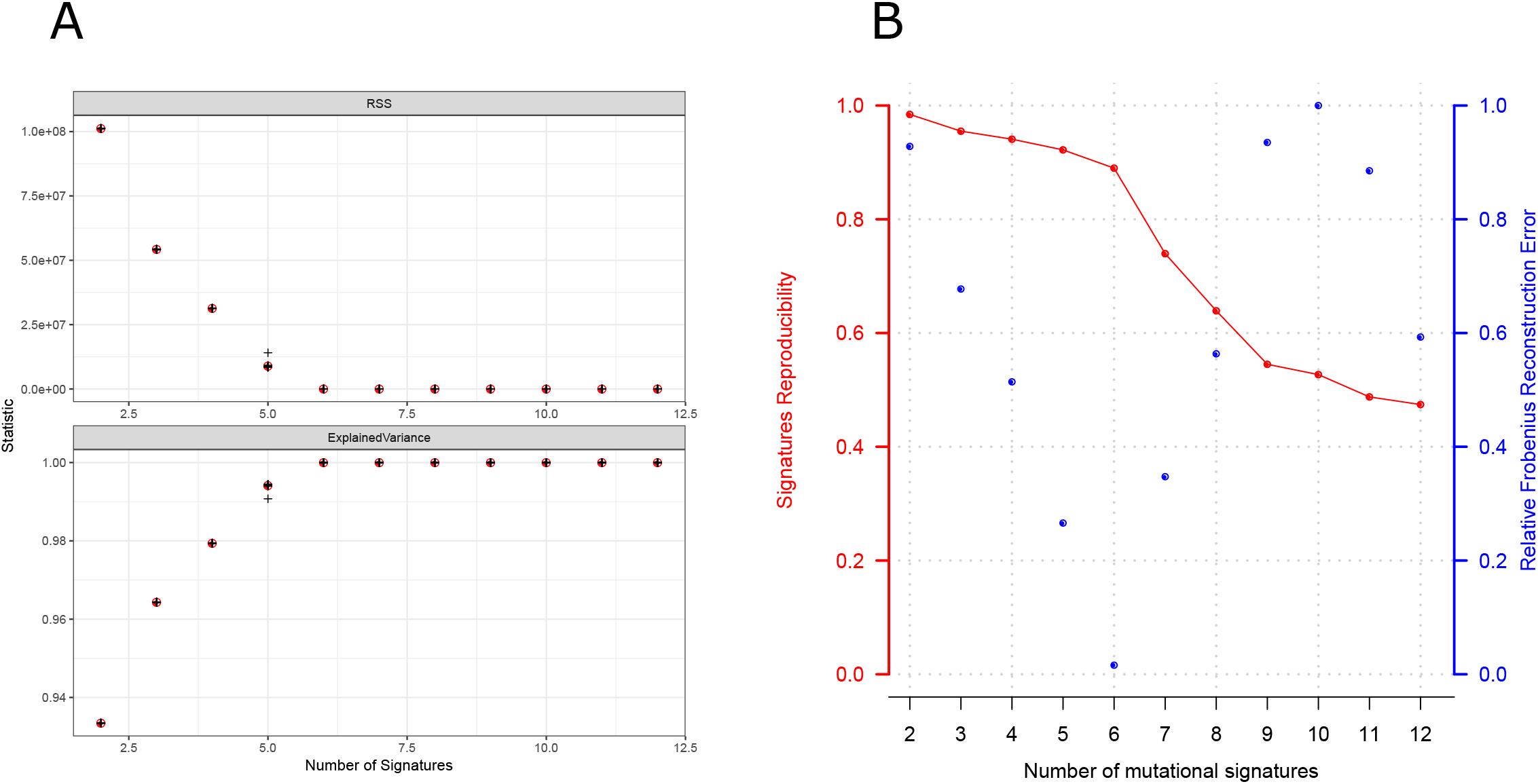
Evaluation diagrams produced by SomaticSignatures and CANCERSIGN packages when applied to the sample dataset (for tool comparison). (**A**) For SomaticSignatures package. (**B**) For CANCERSIGN package.

**Figure 11.**
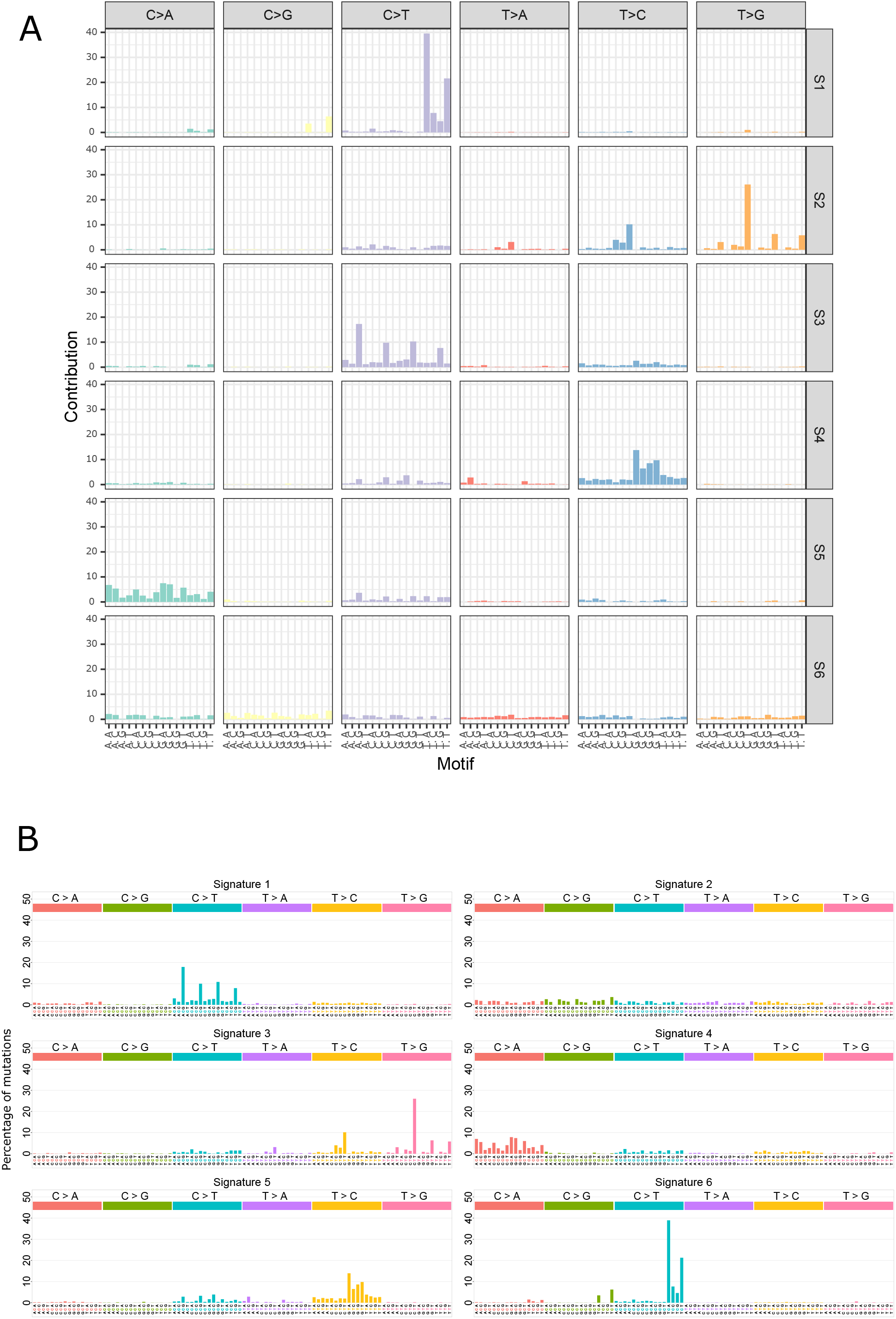
Mutational signatures deciphered by SomaticSignatures and CANCERSIGN packages when applied to the sample dataset (for tool comparison). (**A**) For the SomaticSignatures package. (**B**) For the CANCERSIGN package.

We also compared the tools in terms of time efficiency. Note that the number of replicates for SomaticSignatures is equivalent to the number of bootstraps for CANCERSIGN. It is observed that in the same amount of time (~45 minutes), the number of bootstraps performed by CANCERSIGN was more than the number of replicates in the process of SomaticSignatures (100 vs. 20). Thus, we can conclude that CANCERSIGN performs the analyses faster than SomaticSignatures.

## CONCLUSION

Systematic analyses of mutations in tumor biopsies from a large number of cancer types have identified at least 30 mutational signatures, each pointing to distinct molecular mechanisms acting on cellular DNA. The computational method used to extract mutational signatures is becoming an integral part of cancer research. Therefore, in recent years, a number of studies have focused specifically on the development of bioinformatics tools for analysis of mutational signatures. In the present study, we reported the development of a stand-alone tool called CANCERSIGN, which does not require any programming skills to be used. This tool has several unique features. Firstly, it has been optimized to run parallel computational analysis in order to speed up *de novo* mutational signature extraction. Secondly, it enables the extraction of 5-mer mutational signature profiles in addition to the commonly used 3-mer signature patterns. Thirdly, using the clustering features of this tool, differences between patients and/or tumor types can be investigated using all mutation profiles and/or subsets of selected mutations. In addition, using this tool, we identified a number of novel signatures. Overall, CANCERSIGN is a multi-functional and user-friendly computational tool for accurate and quick analysis of mutation signatures and clustering of tumor samples. This tool is a stand-alone package, is freely available, and does not require any specific computational skills to run.

## AUTHOR CONTRIBUTION

HAR designed the study. MB, HAR, HRR, DE, ARRF wrote and revised the manuscript. HAR, FV provided the data and revised the manuscript. MB, HRR and HAR carried out tool implementation. MB generated all figures and tables. MB, HAR and HRR and MM performed statistical analyses. All authors have read and approved the final version of the paper.

## DATA AVAILIBILITY

All supplementary files are available on the journal website.

## SUPPLEMENTARY DATA

The CANCERSIGN is accessible at https://github.com/ictic-bioinformatics/CANCERSIGN and https://github.com/masroor-bayati/cancer_analysis_tool. The CANCERSIGN source code and tool and instructions on how to run CANCERSIGN are provided at https://github.com/masroor-bayati/cancer_analysis_tool.

## ACKNOWLEDGMENT

We would like to kindly thank Ludmil Alexandrov (Department of Cellular and Molecular Medicine and Department of Bioengineering and Moores Cancer Center, University of California, San Diego) for useful discussion on the signatures and his insightful comments on the paper and CANCERSIGN.

## FUNDING

H.A.R. is a research associate in the Harry Perkins Institute of Medical Research. A.R.R.F. is supported by a Senior Cancer Research Fellowship from the Cancer Research Trust. This work was supported by funds raised by the MACA Ride to Conquer Cancer. This work has also been supported by Iran National Science Foundation (INSF) Grant No. 96006077.

## CONFLICT OF INTEREST

The authors declare no competing financial interests.

## SUPPLEMENTARY FIGURES LEGENDS

**Figure S1|Tool functionality map. This diagram abstracts the functional features of our tool.** The user can either choose to extract mutational signatures in the conventional format which are the spectrums over 96 mutation types (3-mer motifs) or extract 5-mer mutational signatures. The box indicating “NMF algorithm” consists of several steps. Its main steps are bootstrapping from mutational catalogue matrix, NMF iteration and clustering the results obtained from the bootstrapped data. In the clustering part, the user can choose different sets of features to cluster the samples. These feature sets are based on either mutational motifs or mutational signatures. In the former case, counts or proportions of 3-mer or 5-mer mutational motifs can be selected, and in the latter case, contributions of 3-mer or 5-mer mutational signatures can be chosen for clustering features.

**Figure S2|Evaluation plot for deciphering 3-mer mutational signatures.** Whole genome samples in breast cancer.

**Figure S3|Evaluation plot for deciphering 3-mer mutational signatures.** Whole genome samples in blood cancer.

**Figure S4|Evaluation plot for deciphering 3-mer mutational signatures.** Whole genome samples in pancreas cancer.

**Figure S5|Evaluation plot for deciphering 3-mer mutational signatures.** Whole genome samples in stomach cancer.

**Figure S6|Evaluation plot for deciphering 3-mer mutational signatures.** Whole genome samples in brain cancer.

**Figure S7|Evaluation plot for deciphering 3-mer mutational signatures.** Whole genome samples in kidney cancer.

**Figure S8|Evaluation plot for deciphering 3-mer mutational signatures.** Whole genome samples in liver cancer.

**Figure S9|Evaluation plot for deciphering 3-mer mutational signatures.** Whole genome samples in prostate cancer.

**Figure S10|Evaluation plot for deciphering 3-mer mutational signatures.** Whole genome samples in bladder cancer.

**Figure S11|Evaluation plot for deciphering 3-mer mutational signatures.** Whole genome samples in colorectal cancer.

**Figure S12|Evaluation plot for deciphering 3-mer mutational signatures.** Whole genome samples in Oesophagus cancer.

**Figure S13|Evaluation plot for deciphering 3-mer mutational signatures.** Whole genome samples in ovary cancer.

**Figure S14|Evaluation plot for deciphering 3-mer mutational signatures.** Whole genome samples in head & neck cancer.

**Figure S15|Evaluation plot for deciphering 3-mer mutational signatures.** Whole genome samples in skin cancer.

**Figure S16|Evaluation plot for deciphering 3-mer mutational signatures.** Whole genome samples in lung cancer.

**Figure S17|Evaluation plot for deciphering 3-mer mutational signatures.** Whole genome samples in bone cancer.

**Figure S18|Evaluation plot for deciphering 3-mer mutational signatures.** Whole genome samples in uterus cancer.

**Figure S19|Evaluation plot for deciphering 3-mer mutational signatures.** Whole genome samples in nervous system cancer.

**Figure S20|Evaluation plot for deciphering 3-mer mutational signatures.** Mixed data (whole genome and whole exome) samples in breast cancer.

**Figure S21|Evaluation plot for deciphering 3-mer mutational signatures.** Whole exome samples in breast cancer.

**Figure S22|Evaluation plot for deciphering 3-mer mutational signatures.** Mixed data (whole genome and whole exome) samples in skin cancer.

**Figure S23|Evaluation plot for deciphering 3-mer mutational signatures.** Whole exome samples in skin cancer.

**Figure S24|Evaluation plot for deciphering 5-mer mutational signatures.** Whole genome samples in breast cancer.

**Table S1|General statistics of the dataset of somatic mutations.**

**Table S2|Numerical values of deciphered 3-mer mutational signatures.** Whole genome samples in breast cancer.

**Table S3|Numerical values of deciphered 3-mer mutational signatures.** Whole genome samples in blood cancer.

**Table S4|Numerical values of deciphered 3-mer mutational signatures.** Whole genome samples in pancreas cancer.

**Table S5|Numerical values of deciphered 3-mer mutational signatures.** Whole genome samples in stomach cancer.

**Table S6|Numerical values of deciphered 3-mer mutational signatures.** Whole genome samples in brain cancer.

**Table S7|Numerical values of deciphered 3-mer mutational signatures.** Whole genome samples in kidney cancer.

**Table S8|Numerical values of deciphered 3-mer mutational signatures.** Whole genome samples in liver cancer.

**Table S9|Numerical values of deciphered 3-mer mutational signatures.** Whole genome samples in prostate cancer.

**Table S10|Numerical values of deciphered 3-mer mutational signatures.** Whole genome samples in bladder cancer.

**Table S11|Numerical values of deciphered 3-mer mutational signatures.** Whole genome samples in colorectal cancer.

**Table S12|Numerical values of deciphered 3-mer mutational signatures.** Whole genome samples in esophagus cancer.

**Table S13|Numerical values of deciphered 3-mer mutational signatures.** Whole genome samples in ovary cancer.

**Table S14|Numerical values of deciphered 3-mer mutational signatures.** Whole genome samples in head & neck cancer.

**Table S15|Numerical values of deciphered 3-mer mutational signatures.** Whole genome samples in skin cancer.

**Table S16|Numerical values of deciphered 3-mer mutational signatures.** Whole genome samples in lung cancer.

**Table S17|Numerical values of deciphered 3-mer mutational signatures.** Whole genome samples in bone cancer.

**Table S18|Numerical values of deciphered 3-mer mutational signatures.** Whole genome samples in uterus cancer.

**Table S19|Numerical values of deciphered 3-mer mutational signatures.** Whole genome samples in nervous system cancer.

**Table S20|Numerical values of correlation of 77 signatures identified by CANCERSIGN with 30 previously reported signatures by Nik-Zainal et al. [15].**

**Table S21|Numerical values of deciphered 5-mer mutational signatures.** Whole genome samples in breast cancer.

